# Rosetta’s Predictive Ability for Low-Affinity Ligand Binding in Fragment-Based Drug Discovery

**DOI:** 10.1101/2022.10.18.512794

**Authors:** Elleansar Okwei, Shannon T. Smith, Brian J. Bender, Brittany Allison, Soumya Ganguly, Alexander Geanes, Xuan Zhang, Kaitlyn Ledwitch, Jens Meiler

**Affiliations:** Department of Chemistry, Vanderbilt University, Nashville, TN 37235, USA; Center for Structural Biology, Vanderbilt University, Nashville, TN 37240, USA; Program in Chemical and Physical Biology, Vanderbilt University, Nashville, TN 37240, USA; Department of Pharmacology, Vanderbilt University, Nashville, TN, 37240, USA; Institute of Drug Discovery, Faculty of Medicine, University of Leipzig, 04103 Leipzig, Germany

**Keywords:** Protein-small molecule binding, SOFAST-HMQC, NMR, HisF, computational docking, Rosetta, RosettaLigand, Surflex

## Abstract

Fragment-based drug discovery begins with the identification of small molecules with a molecular weight of usually less than 250 Da that weakly bind to the protein of interest. This technique is challenging for computational docking methods as binding is determined by only a few specific interactions. Inaccuracies in the energy function or slight deviations in the docking pose can lead to the prediction of incorrect binding or difficulties in ranking fragments in *in silico* screening. Here we test RosettaLigand by docking a series of fragments to a cysteine-depleted variant of the TIM-barrel protein, HisF. We compare the computational results with experimental NMR spectroscopy screens. NMR spectroscopy gives details on binding affinities of individual ligands, which allows assessment of the ligand-ranking ability by RosettaLigand, and also provides feedback on the location of the binding pocket, which serves as a reliable test of RosettaLigand’s ability to identify plausible binding poses. From a library screen of 3456 fragments, we identified a set of 31 ligands with intrinsic affinities to HisF with dissociation constants as low as 400 µM. The same library of fragments was blindly screened *in silico*. RosettaLigand was able to rank binders before non-binders with an area under the curve (AUC) of the receiver operating characteristics (ROC) of 0.74. The docking poses observed for binders agreed with the binding pocket identified by NMR chemical shift perturbations for all fragments. Taken together, these results provide a baseline performance of RosettaLigand in a fragment-based drug discovery setting.

## Introduction

### Computational techniques serve to accelerate the drug discovery and lead development process

Predicting the binding of drug compounds to target structural models using computational techniques is one way to accelerate the drug discovery and lead development processes. As such, various computational algorithms developed over the past few decades have been dedicated to this problem^*1*^. The two main approaches to these computational predictions are ligand-based and structure-based computer-assisted drug discovery (LB- and SB-CADD). In LB-CADD, cheminformatics is applied to a set of ligands of known activity to attempt to describe features that distinguish actives from non-actives such as substructure scaffold, hydrogen bond donors, and molecular weight^*2-4*^. As this method is solely ligand-based, LB-CADD requires no knowledge of the structure of the target protein. On the other hand, SB-CADD uses the structure of the target protein to guide docking scenarios that can estimate degree of fit for various ligands and attempt to predict binding energy^*5, 6*^. Both strategies have proven useful in the field of drug discovery and new developments continue to improve upon their predictive abilities through rigorous testing.

### SB-CADD can rapidly advance initial hit discovery

In SB-CADD, ligands are docked into a known structure of the target protein in a process called virtual high throughput screening. The goal of this strategy is to identify hits (i.e., ligands that bind in the available pocket). Since the goal is to rapidly screen a large library of compounds, the energy function is designed for speed and often uses rough estimations of physical properties to improve the throughput in finite time^*7, 8*^. These energy functions allow for rapid sampling of conformational space of the protein-ligand complex, and often result in near-native predicted binding poses.

As these scoring functions are enhanced for increased sampling capability, the resulting scoring function cannot consider the higher levels of theory required for accurate energy predictions upon binding, and therefore struggles to correctly predict the binding affinity of a complex. Although some interaction types such as hydrogen bonding and certain Van der Waals interactions are well-characterized implicitly or explicitly, in the case of others such as long-range polar interactions from second or third shell effects and desolvation penalties upon ligand binding, it is more difficult to predict energetic effects without quantum-level calculations. As a result, a commonly used strategy for SB-CADD predictions is to use one program for sampling potential protein-ligand interactions and a second, higher level calculation to more accurately score compounds^*9-11*^.

### RosettaLigand protocol uses enhanced sampling method followed by higher-resolution scoring

Rosetta, a macromolecular modeling software suite for protein structure prediction and design^*12, 13*^, has two energy functions that are optimized separately for low-resolution sampling and high-resolution sampling with scoring. The low-resolution scoring function has been optimized for rapid sampling of both translational and rotational ligand changes and uses a grid searching method to increase computational speed. This energy function primarily calculates steric-based interactions while using a “softer” repulsive term, allowing for sampling close to protein atoms without large energetic penalties. These potential clashes can then be resolved during the high-resolution sampling stage where the repulsive term has been ramped to mimic a full Van der Waals term. ^*5, 14, 15*^. Further, RosettaLigand is capable of fully flexible docking in which both the ligand and protein are allowed to sample various conformations to allow for induced fit binding. This strategy has been leveraged successfully in the docking predictions of ligands for crystal structures^*16*^, comparative models^*17-21*^, and even in the case of enzyme design^*22-25*^, which relies on extremely accurate detailing of ligand-protein contacts.

### Fragment-Based Drug Discovery Identifies Hits Using a Focused Library

A common theme in the above studies is the use of drug compounds with high potency and several binding interactions between the drug and protein. Identification of drug compounds via standard high throughput screening requires hundreds of thousands of compounds in a library to cover the broad chemical space of compounds of this size (∼500 Da). An alternative approach to screening drug-like compounds is to use fragment-based drug discovery (FB-DD) which uses smaller compounds (average molecular weight of 200 Da) that provide starting hits for subsequent lead development^*26*^. Multiple hit fragments can subsequently be combined to increase the affinity of these compounds^*27*^. By screening substructures, these libraries only need to be on the order of several thousand compounds^*28*^.

### NMR allows for simultaneous determination of binding affinity and interaction site to aid

*FB-DD* In FB-DD, the small size of the fragments results in few specific binding interactions between the fragment and the target protein. As such, the binding affinities are fairly weak and millimolar affinities are considered reasonable starting points^*29*^. Biophysical techniques such as nuclear magnetic resonance (NMR) spectroscopy, isothermal titration calorimetry (ITC), and surface plasmon resonance (SPR) that are sensitive to weak binding interactions are needed to screen a fragment library. NMR spectroscopy has proven extremely useful in this regard as this technique provides a measure of low-affinity binding while also identifying the binding pose to the protein target^*30*^. Further, the use of SOFAST-HMQC allows for rapid screening by NMR^*31*^.

### Testing RosettaLigand’s ability for in silico FB-DD

Here, we benchmarked the predictive ability of Rosetta^*13, 32*^ to accurately dock fragments to a protein target. This was particularly challenging for the docking algorithm as these fragments possessed low affinity and needed only a few interactions to bind the protein. For this test, we identified a set of fragments that bound to a TIM-barrel protein, HisF, with affinities in the high micromolar range, via NMR chemical shift analysis. These fragments were blindly docked into the binding pocket of HisF, in which we determined the X-ray crystallography structure. This docking strategy was successful at distinguishing actives from inactives, which is indicative of a discriminatory scoring function. Further, we found that the docking poses of the actives, guided only by Rosetta’s sampling algorithm, correlated strongly with the residues identified in the NMR experiments.

We wanted to assess the quality of docking predictions as if there was no previous structural knowledge of the orientation of known binders. Typically, in a drug discovery pipeline, structural information of existing binders is used, alignments of libraries are performed on known binders, and predictions are made from this dataset. Although we chose a use-case that had a co-crystal structure of a bound ligand, we wanted to know if these docking predictions yielded similar results to an alignment-based approach. To investigate this, a ligand-based prediction was performed using SurflexSim^*33*^ to align these compounds with the previously crystallized native ligand for HisF. It was observed that output docking poses matched closely to Rosetta’s predicted binding pose. Taken together, these results show that Rosetta can reliably dock and score fragments despite their low affinity and number of specific interactions. This suggests that Rosetta is a valid starting tool for FB-DD and subsequent drug design.

## Methods

### HisF Mutagenesis, Expression, and Purification

The sequence of wild-type HisF (HisF^wt^), obtained from PDB entry 1THF, was encoded in the pBG100 vector with an N-terminal hexahistidine purification tag and TEV protease cleavage site. HisF^C9S^ mutant was generated using QuikChange PCR with the primers HisF^C9S^-Fwd 5’-CTGGCGAAGCGTATTATCGCGAGCCTGGACGTTAAAGACGGTCGC-3’ and HisF^C9S^-Rev 5’-GCGACCGTCTTTAACGTCCAGGCTCGCGATAATACGCTTCGC CAG-3’. The plasmids encoding WT HisF or HisF-C9S were transformed into BL21(DE3) cells for bacterial expression. Small-scale cultures were grown overnight in LB media. The cells were centrifuged and the pellets transferred into M9 media containing 0.5 g/L (^15^NH_4_)_2_SO_4_ (Cambridge Isotopes) and grown at 37°C to an OD_600_ of 0.5. Growth cultures were then moved to room temperature and incubated with shaking until an OD_600_ of 0.6 – 0.8 before inducing protein expression with IPTG for overnight growth. The cells were harvested by centrifugation at 6500 rpm for 20 minutes, and stored at -80°C. Cell pellets were lysed in 20 mM Tris buffer, 5 mM imidazole, 200 µg/mL lysozyme and Roche EDTA-free protease tablets at pH 7 followed by sonication. The clarified soluble fraction was purified over packed TALON resin and the immobilized protein was eluted with excess imidazole (250 mM). The protein was dialyzed against 10 mM MES, 50 mM KCl, and 1 mM EDTA at pH 6.8 and stored at -30°C.

#### Crystallization of HisF^C9S^

HisF^C9S^ was dialyzed against 25 mM potassium phosphate and 150 mM NaCl at pH 7.5. Crystals of HisF^C9S^ were grown using vapor diffusion. HisF^C9S^ was concentrated to 10 mg/mL and set in 24-well hanging drop plates using a ratio of 1:1 with well solution. Final crystals were obtained in a solution of 100 mM Tris and 25% PEG-3350 at pH 7.5. Crystals were flash frozen after briefly soaking in mother liquor containing 25% ethylene glycol. X-ray diffraction was carried out at low temperatures on an in-house Bruker Microstar instrument, with the crystal diffracting to 1.9 Å. The data was phased using Phaser on an ensemble of templates (1THF, 2AON, 1VH7) generated with Ensembler. Phenix.Refine was used for additional refinement.

### NMR data collection, analysis, and backbone assignment transfer

NMR spectra were collected on uniformly labeled ^15^N-HisF^wt^ and ^15^N-HisF^C9S^ at a concentration of 100 µM. All samples were spiked with 5% D_2_O to lock the signal. Experiments were performed on a 600 MHz Bruker spectrometer at 25°C equipped with a QCI-P cryoprobe and a sample jet. Backbone assignments of HisF^wt^ were transferred from a previously published ^1^H-^15^N TROSY-HSQC spectrum (BMRB: 15741) to our ^1^H-^15^N HSQC spectra^*34*^. The assignments of HisF^wt^ were then transferred to the ^1^H-^15^N SOFAST^*35*^ HMQC spectra of HisF^C9S^ and were used for successive rounds of small molecule screenings and titrations. All NMR spectra were processed using NMR pipe^*36*^ and analyzed with Sparky^*37*^.

### Small Molecule Screening

The Vanderbilt fragment library was screened using a 96-well plate NMR setup to identify weak binders of HisF^C9S^. Three plates consisting of twelve compounds per well were screened to identify candidate molecules. The individual concentration of the twelve compounds in each well was 600 µM. The mixture of twelve compounds in each well was dissolved with 100µM HisF^C9S^ protein in NMR buffer with 4% DMSO and 10% D_2_O. Small molecules were screened by observing changes in ^1^H-^15^N backbone chemical shifts of HisF^C9S^ residues near the binding pocket during SOFAST HMQC experiments. Wells displaying chemical shifts were partially de-convoluted by screening ligands in groups of four. Ligands in wells that displayed chemical shifts were then screened individually to identify the compound(s) that exhibited these effects. Hits identified in this screen were compared to the remaining compounds in the Vanderbilt fragment library using Chemcart. The search was filtered by atomic charge, bond type, and match primary fragment and was limited to small molecules with 0 – 3 rotatable bonds as this was the range of the search set. A total of 86 related fragments were identified and screened against HisF^C9S^.

### NMR Ligand Titration to Determine Binding Affinity

Hits identified during small molecule screening were further investigated to experimentally determine the binding affinity. An excess of up to 12 molar equivalent of small molecule was titrated to 100 µM HisF^C9S^ and the ^1^H-^15^N chemical shift perturbations were monitored through SOFAST-HMQC experiments^*35*^. Combined ^1^H and ^15^N chemical shifts of the perturbed peaks were calculated using Equation 1^*38, 39*^.

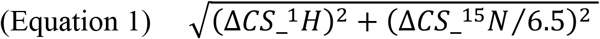

Here, Δ*CS_*^1^*H* is the chemical shift difference between small molecule bound and unbound protein for amide proton and Δ*CS_*^15^*N* for amide nitrogen. Concentration-response curves (CRC) were prepared in GraphPad Prism (GraphPad Prism version 6.04 for Windows, GraphPad Software, La Jolla California USA, www.graphpad.com)^*40*^.

### RosettaLigand Blind Docking of Fragments

The fragments were docked into a model of HisF^C9S^ using RosettaLigand as described previously^*19, 41*^. Mol files of the fragments were used to generate conformers using confab. The output conformer mol files were then converted into Rosetta-readable params and pdb files. Since the binding site was unknown, fragments were placed in the center of the large convex face of HisF^C9S^ using the StartFrom mover in RosettaScripts^*42*^. The entire width of this pocket (10 Å) was used as the potential binding region and defined the box size (radius) as 6 Å such that the scoring grid would cover a space just larger than the potential binding pocket. In the first round of docking, the fragments were allowed freedoms of 1 Å step sizes and full 360° rotation in the Transform mover^*5*^. 5000 models were generated in the first round of docking. The models were sorted by total Rosetta energy unit (REU) score and then the top 10% were sorted by interface energy. The top 50 models by interface energy were used as input in the second round of docking. In the second round of docking, fragment movement was reduced to 0.2 Å step sizes and 45° rotation in the Transform mover. Again 5000 models were generated (100 from each parent model), and the same sorting scheme was used to identify the top 50 models. To ensure model diversity, no more than 4 models from a single parent model (or less than 10% of the 50 models) were allowed in the final set of 50 models. These 50 models were then used in a last round of docking in which the fragment freedom was reduced further to 0.04 Å step sizes and 5° rotation in the Transform mover. 5000 models were generated. After selecting for the top 10% by REU score and sorting by interface score, the top 10 models were selected for analysis. In every stage of docking, the InterfaceScoreCalculator^*17*^ was used to determine the interface score of the fragment-protein complex. This is calculated by scoring the model of the complex and then moving the fragment 1000 Å away from the protein and rescoring.

### Assessment of RosettaLigand’s Discrimination of Actives from Inactives

The full set of experimentally tested drug fragments were docked in the binding pocket of HisF^C9S^ as described above and ranked. These rankings were compared to the list of fragments that displayed binding interactions. A receiver-operator curve (ROC) was generated using R analysis.

### ROC Analysis of Rosetta Binding Site Predictions

Rosetta binding site predictions were analyzed using the ddg mover in RosettaScripts^*42*^ (see Supplemental Methods). Energies of the bound protein-fragment complex were compared to those of the protein *apo* state and differences in the energy of the two states were computed on a per residue level. The energies of the top ten binding poses were averaged. Averages that displayed a non-zero energy between the two states were assigned as Rosetta-predicted binding sites and the magnitude of the energy was used to rank the strength of the predicted residues. Binding pocket residues were identified as those that displayed a displacement of the ^1^H-^15^N chemical shift upon titration of the fragment. The accuracy of the predicted binding site residues was determined using the ROCR library in R^*43*^. A modification to this analysis was that for residues that displayed a displacement in the HSQC spectrum, Rosetta energies were queried for that given residue (i) and the residues adjacent (i–1 and i+1) for non-zero values as the displacement of a backbone-amide chemical shift may result from ligand interaction with any of these three residues. Measures were calculated as a true positive rate compared to a false positive rate and the area under the curve was determined. Only those residues which have been assigned in the HSQC spectrum (63 of 253 residues) were used for this analysis to not over-calculate the true negative rate.

### Surflex-Sim Alignments to Binding Mode Hypotheses

Surflex-Sim as implemented in the SybylX 2.1.1 suite of computational chemistry programs was used to generate alignments of HisF binders and non-binders to either computed binding mode hypotheses or the PRFAR crystal structure^*2*^. The PRFAR structure was trimmed to exclude the flexible sidechain past the amidine moiety to constrain alignments to a physically realistic region of space. For these alignments, ring flexibility was considered, pre- and post-run energy minimizations were performed, spin density was set to 5, number of spins per molecule was set to 20, and up to 50 conformations of each molecular fragment were investigated. This choice of parameters was made to ensure that a thorough sampling of both conformational and 3-dimensional search space was performed when aligning compounds. Once completed, the most similar conformation of each compound to the hypothesis (according to score) was extracted and used for analysis.

## Results

### Selection of HisF^C9S^ for NMR and Computational Experiments

*We selected the (βα)8-barrel (TIM-barrel) protein HisF from Thermatoga maritima as a model system for these studies due to the availability of both the NMR and X-ray experimentally-determined structures. We obtained the protein sequence of HisF from the PDB entry 1THF*^*44*^, *the highest resolution structure of HisF. Of note, the protein sequence from 1THF contains a Thr21Ser point mutation. As this mutation is conservative and outside of the ligand binding pocket, we consider this our wild-type HisF sequence. Upon analysis of the ligand binding pocket prior to screening, we identified a cysteine at position 9 which we mutated to serine to avoid the potential of covalent interactions between fragments and the protein. The X-ray crystal structure for the HisF*^*C9S*^ *mutant is identical to HisF*^*wt*^ *and was determined at a 1*.*9 Å resolution (PDB 5TQL*,). The full-atom RMSD between HisF^C9S^ and 1THF is 0.85 Å (Figure 1A). Further, the Ser9 sidechain adopts the same rotameric state in HisF^C9S^ as Cys9 in 1THF. Taken together, we conclude that the experimentally-determined HisF^C9S^ structure is nearly identical to HisF^wt^ and provides a suitable model protein for NMR screening and docking studies.

**Figure 1.**
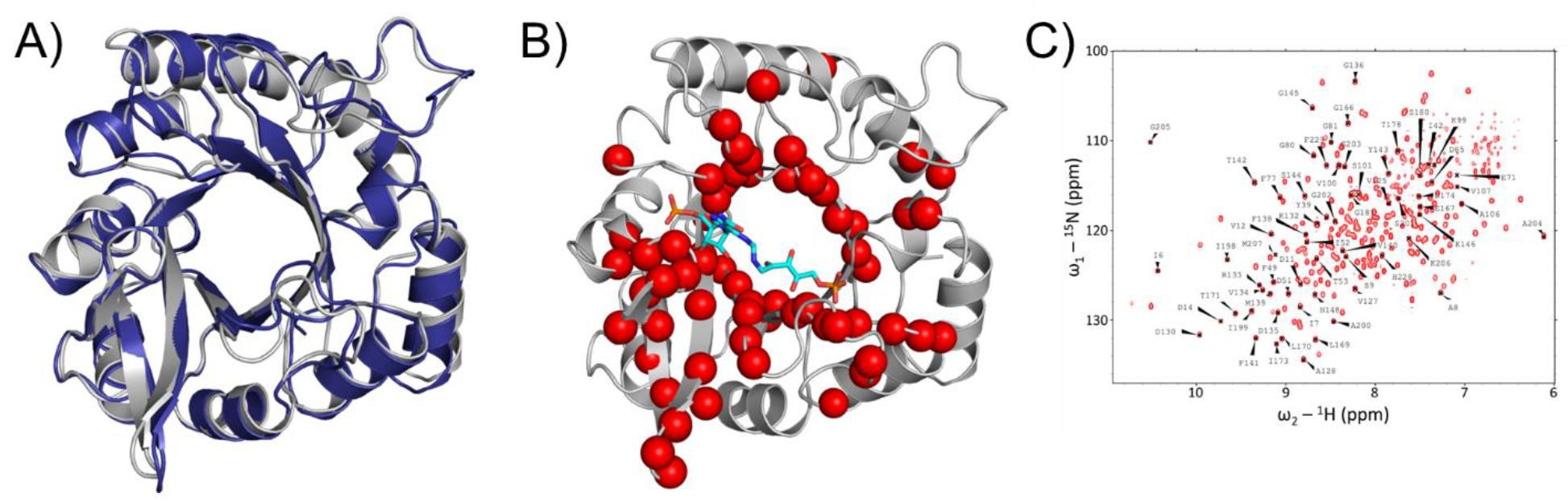
The HisF binding pocket. (**A**) Comparison of the *Thermatoga maritima* wild-type HisF crystal structure in blue (PDB ID: 1THF) with HisF^C9S^ in grey (PDB ID: 5TQL). The superimposed structural models have an RMSD of 0.853 Å. (**B**) After transferring the NMR assignments from HisF^wt^ to HisF^C9S^, a subset of backbone amides were selected based on their proximity to the binding pocket as inferred from the yeast HisF-PRFAR complex (PDB ID: 1OX5). PRFAR from the crystal complex is shown in cyan. The selected backbone amide groups are shown as red spheres. (**C**) ^1^H-^15^N HSQC-TROSY of HisF^C9S^. Tracked amide resonances from panel B are marked and labeled.

### NMR assignment of HisF^C9S^ spectra

Amide ^1^H-^15^N resonances from a previously assigned HisF spectrum^*34*^ collected at 30°C were transferred to a ^1^H-^15^N HMQC spectrum we obtained of HisF^wt^ at 25°C. We subsequently generated a ^1^H-^15^N SOFAST-HMQC of HisF^C9S^ at 25°C and transferred assignments for 150 residues from our HisF^wt^ spectrum. Since the goal of this study is to identify small molecules that bind within the native ligand site, attention was focused on the known binding pocket of HisF for its natural substrate 5-[(5-phospho-1-deoxyribulos-1-ylamino)methylideneamino]-1-(5- phosphoribosyl)imidazole-4-carboxamide (PRFAR). Of the 150 confirmed residues, an inclusive list of 63 residues was compiled to be within or proximal to the binding pocket, based on residues that were confidently assigned within two shells of the yeast HisF-PRFAR binding pocket and were tracked over the remaining experiments (Figure 1B + C).

### Identification of Potential Hits to HisF^C9S^

The Vanderbilt fragment library of around 14,000 fragments was built from drug-like fragments that contain substructures that often bind proteins such as carboxylic acids and heterocycles^*45*^ and generally adhere to the ‘rule of three’ meaning they have a molecular weight less than 300 Da, a ClogP less than 3, and contain less than 3 hydrogen bond acceptors^*46*^. 3456 fragments from the library were screened against HisF^C9S^ to identify fragments with intrinsic binding affinity. Binding of fragments were tested using the ^15^N-SOFAST heteronuclear multiple quantum correlation (SOFAST-HMQC) NMR technique, which allows for rapid detection of conformational changes at the binding interface induced by small molecule binding^*31*^. The 3456 fragments were batched on 96-well plates with 12 fragments per well^*30, 31*^. Fragments in wells that displayed chemical shift perturbations were tested individually against HisF^C9S^ for binding activity. A set of 25 hits were identified in this screen. The remaining ∼10,000 compounds in the Vanderbilt fragment library were searched for similar fragments using substructure similarity as implemented in the Chemcart software. A total of 86 related fragments were screened against HisF^C9S^ and 15 were identified as additional hits. Combined with the initial 25 confirmed hits, a total of 40 fragments were discovered that possessed intrinsic binding affinity for HisF^C9S^.

### Binding Analysis of Individual Hits

The 40 fragments identified as hits were confirmed by determining the binding affinity through NMR titration experiments. Chemical shift perturbations were monitored in HisF^C9S^ in the presence of each fragment at concentrations up to 1200 µM. A combined ^1^H and ^15^N chemical shift difference was calculated against the apo NMR spectrum according to Equation 1^*38, 39*^. This value was plotted against fragment concentration to derive binding curves. Figure 2 highlights the titration spectra and binding curve of one of these ligands. Out of the 40 titration experiments, only 31 hits could be fit with a one-site binding model (Supplementary Figure 1). The 9 remaining hits all generated chemical shift perturbations during titration experiments but could not be fit to a simple one-site binding model and were removed from the dataset. The K_D_ values of these naïve binders are, as expected, weak with the best affinities approaching 400 µM. In many instances, K_D_ values are affiliated with a substantial error as the binding is not saturated at the highest measured fragment concentration.

**Figure 2.**
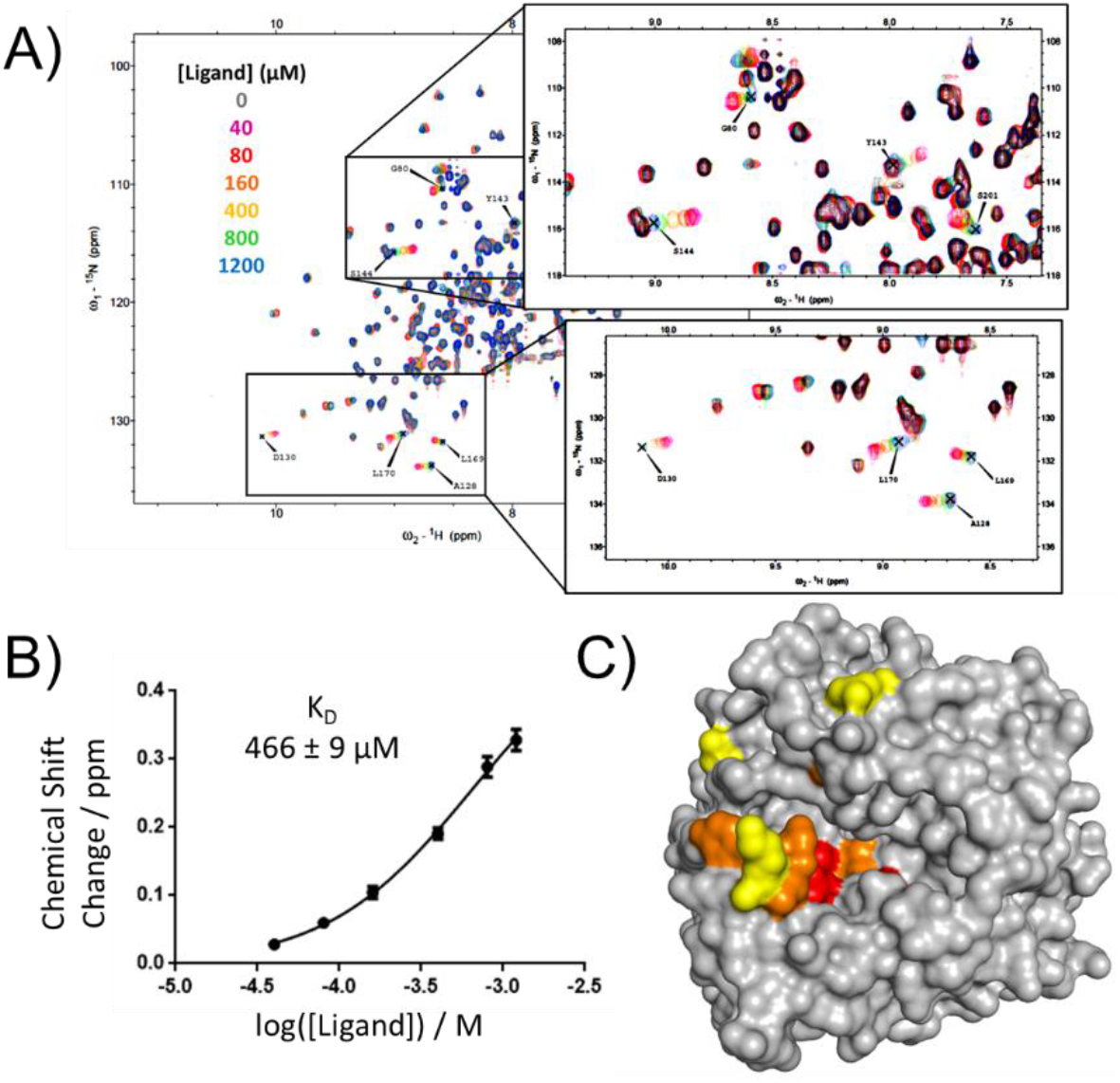
Identification of Low-Affinity Ligands for HisF^C9S^. (**A**) An example NMR titration of compound VU0139210. HMQC-NMR spectra of HisF^C9S^ in the presence of 0, 40, 80, 160, 400, 800, and 1200 µM ligand reveal peaks that shift in the presence of increasing ligand concentration. Zoomed in inserts show tracked peaks that exhibit the largest chemical shifts. (**B**) The plotted binding curve of chemical shift change (ppm) vs log ligand concentration (M) gives a K_D_ 466 µM. (**C**) Residues that were identified as responding to ligand binding across the full set of 31 low-affinity ligands are mapped onto the protein surface. The residues are colored according to frequency of chemical shift perturbation throughout the ligand series with yellow-orange-red representing lowest to highest frequency, respectively.

### Binding Mode of the Small Molecules

We analyzed the 31 naïve binders NMR titration data and identified a HisF^C9S^ binding pocket hotspot from the chemical shift perturbations (Figure 2C). Among the 63 tracked resonances, 35 never displayed any peak shifts and these residues, are therefore categorized as not interacting with the fragments. Hot spots for interaction include the catalytic residue Asp130 and many surrounding residues including the nearby loop (residues 142 – 144). Interestingly, catalytic residue Asp11 displayed no peak shifts. According to the model of yeast HisF-PRFAR complex^*47*^, Asp11 interacts with the glycerol phosphate while Asp130 binds the imidazole-ribose region. The binding hotspots for these identified binding fragments cluster around the region that interacts with PRFAR’s aromatic ring moieties which may provide insight into why many of the novel ligands contain aromatic rings.

### RosettaLigand Can Discriminate Active from Inactive Drug Fragments

The full set of 3542 fragments that were screened experimentally were also screened *in silico* against the HisF^C9S^ binding pocket using RosettaLigand. The binding cavity of HisF^C9S^ is quite large with dimensions of 10 Å x 10 Å x 15 Å. The fragments however are mostly planar with a maximum dimension of around 5–8 Å. To test Rosetta’s ability to predict the region in space that the fragments bind to we placed the fragments in the geometric center of the binding pocket prior to sampling of the space^*19, 41*^. A set of 1000 docked poses were generated for each fragment. The fragments were ranked using the predicted binding energy of the top 5% of these poses. The rank was compared against the set of actives and a receiver-operator characteristic (ROC) curve was generated to evaluate true positives from false positives. From this analysis, it was found that the AUC= 0.74, which is on par with many structure-based screening methods^*48*^. We also investigated early enrichment of hits by calculating the logAUC of the top 10% of compounds by score. The logAUC of our set was 0.09 which suggests that this method does not perform well when selecting true actives in the top tier of compounds by ranking. Despite the problem of early enrichment, RosettaLigand, a structure-based docking strategy, can accurately discriminate active from inactive fragments (Figure 3A).

**Figure 3.**
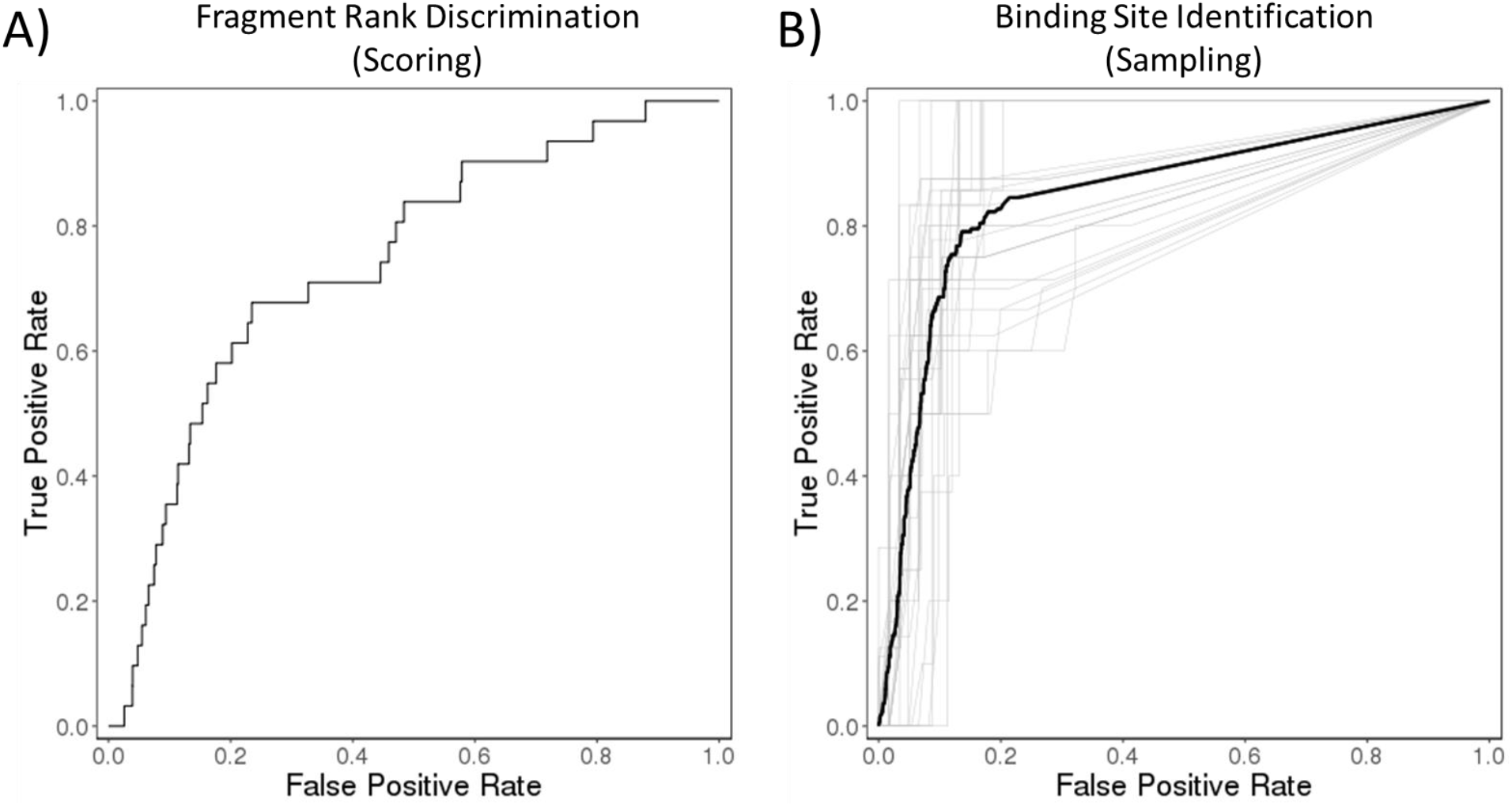
ROC Analysis of Rosetta Predictions. (**A**) Discrimination of actives from inactives using the rank order from average binding energies of all 3546 drug fragments screened by NMR. (**B**) Shown is the receiver-operator curve for Rosetta predictions of residues involved in ligand binding as verified by perturbations to the chemical shift in fragment titration HSQC spectra. The individual fragment analyses are shown in light grey and the aggregate analysis is shown in dark black. We determined an aggregate ROC curve for all 31 ligands with an area under the curve of 0.85.

### High-Density Docking of Active Compounds with RosettaLigand

The 31 confirmed binding fragments were docked into the binding cavity of HisF^C9S^ over multiple rounds to ensure high-density sampling. A round of 5000 docking trajectories with large fragment movements were carried out to allow sampling over the entire pocket (Supplementary Figure 2A). Top models were ranked by binding energy and overall energy. The binding poses were used as starting positions for a subsequent round of docking with tighter restrictions on ligand movement (Supplementary Figure 2B). A last round of docking was carried out on top models again until a consensus binding site was identified (Supplementary Figure 2C). Through each round of docking, the binding mode converged and the energies for the overall complex and binding site improved (Supplementary Figure 2D).

### Structure-Based Fragment Docking Captures Experimentally Determined Binding Pocket

Following the last round of docking, a set of ten models were selected based on binding energy to best represent the likely binding pose. These binding poses localize to the identified binding pocket in HisF^C9S^ as evidenced by chemical shift perturbations. To quantify the degree of agreement between Rosetta predictions and experimental results we used ROC analysis to measure a true positive rate of identification. Per-residue ΔΔGs were calculated over the Rosetta predicted protein-ligand complexes and residues involved in ligand binding were assigned non-zero energies. These residues were compared against the list of chemical shifts that were shifted upon ligand titration. Importantly, if a residue was seen to undergo a movement in the HSQC titration, the Rosetta-determined energies were queried on the i, i+1, and i-1 residues for predicted involvement in ligand binding. This is due to the fact that a backbone-amide chemical shift reports on changes in the chemical environment which can result from local and non-local changes^*49*^. Using this approach, we determined an aggregate ROC curve for all 31 ligands with an area under curve of 0.85 (Figure 3B). For each of the individual ligands, ROC analysis resulted in area under curve values of 0.72-0.98 suggesting that RosettaLigand strongly predicts the binding location of the fragments within the HisF^C9S^ binding pocket.

### Superimposition of HisF^C9S^ Naïve Binders with HisF/substrate Complex

To gain more geometrical insight into trends between chemical structure and binding energy, the crystal structure of PRFAR from the complex with yeast HisF was used to flexibly align all 31 binding fragments using the Surflex-Sim program^*2*^. Since the NMR and Rosetta docking experiments show that these ligands all bind in the same region of the ribose subunit of PRFAR, the flexible portion of PRFAR was removed during these alignments to ensure that physically realistic alignments were made. Eight of the ten strongest binding fragments adopt a conformation orienting polar or negatively charged groups with the phosphate group of PRFAR and the remainder of the ligand aligns with a portion of the ribose or imidazole subunits depending on the molecular size and character. As seen in Figure 4, the overall binding hypothesis is quite similar between the two methods. The use of independent methods that result in similar solutions lends credence to both methods and strengthens our binding mode hypotheses.

**Figure 4.**
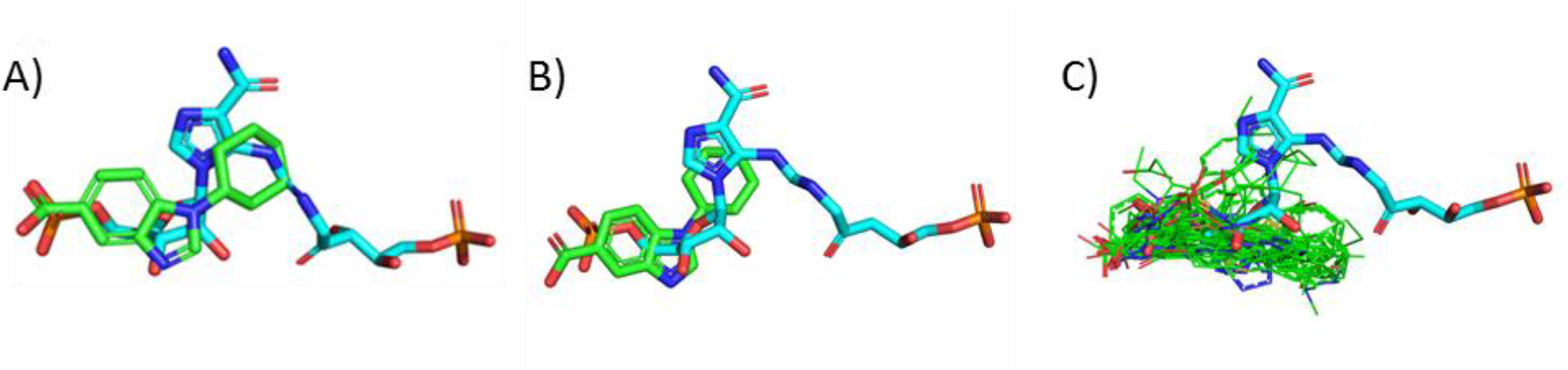
Binding hypotheses by computational modeling. (**A**) The major alignment solution from Surflex-Sim between the HisF naïve binders (green) and the PRFAR crystal structure (cyan). Shown here is VU0151854. (**B**) The binding hypothesis of VU0151854 from RosettaLigand docking again with the ligand in green and PRFAR in cyan. (**C**) The docking solutions of all 31 ligands from RosettaLigand (green) with PRFAR (green) highlight the overall similarity of docked solutions.

## Discussion

### NMR Screening Identifies a Low-Affinity Ligand Dataset for HisF^C9S^

Whereas most tests of ligand docking predictions are performed on high-affinity ligands, we sought to test RosettaLigand’s predictive ability on moderate to low-affinity ligands. We chose HisF as a model system for these studies due to the availability of experimentally-derived structural models from NMR and X-ray crystallography. In this work, our high-resolution crystal structure of HisF^C9S^ provides a reasonable structure for docking studies. Additionally, NMR has the unique advantage to identify low-affinity binding fragments and their binding location(s). The weak affinity of these fragments are too low for co-crystallization. Unlike NMR, other methods of screening such as isothermal calorimetry or surface plasmon resonance would not provide information regarding residues involved in ligand binding. Using NMR screening we were able to generate a dataset of low-affinity binding fragments to HisF^C9S^. After fragment identification, our NMR titration experiments showed that ∼25% of these ligands could not be fit to a one-site binding model. We removed these ligands due to the potential of two or more binding sites, cooperative binding, or non-specific binding.

### RosettaLigand accurately predicts binding of low-affinity fragments

The identification of this dataset provided an ideal opportunity to test the ability of RosettaLigand to predict the binding of low-affinity fragments. Small molecule docking is dependent on highly specific small molecule and side-chain positioning. Accurate prediction of binding for a small molecule fragment is a stringent test of the computational algorithm, as only a few interactions are needed to confer the low affinity. This test was additionally made difficult as the ligands were placed in the geometric center of the binding surface and allowed to dock blindly. As demonstrated in the NMR experiments, the binding site of these ligands was away from the center in a small region. The rank order of ligands is a task generally used in ligand-based drug screening. However, it was seen that even with low sampling RosettaLigand could accurately rank actives from inactives. The limitation here is seen in the lack of early enrichment, as seen by the logAUC value. This is typical of docking-based screening, where although these protocols can sufficiently rank binders favorably, this method is typically not sensitive enough to rank these within the <1% of compounds screened. On the other hand, binding pocket analysis measures the contribution of individual amino acids to the binding of the small molecule as opposed to total binding energy. Comparison of these predictions with the HMQC-NMR chemical shift mapping of residues in the entire binding surface of HisF showed that, despite these difficulties, RosettaLigand showed a strong predictive ability to place these molecules in their respective binding pockets.

### Structure-Based Ligand Docking Results in Similar Ligand Orientation as Ligand-Based Results

The yeast HisF protein has previously been crystalized with the native ligand PRFAR. Despite large differences between the identified ligand and PRFAR chemical structure, rough similarities such as a negatively charged group linked to an aromatic moiety exist. Alignment of these groups between the novel ligands and PRFAR would provide an ideal starting point for docking studies. As seen in the Surflex-Sim results, such an alignment provides highly similar binding poses despite no input regarding the protein structure. The ability to dock the ligands as a group or with alignment to a known ligand would likely have decreased the number of decoys needed to identify accurate binding poses^*50*^. The similarity between the Surflex-Sim aligned binding poses and the correlation with experimental results further strengthen the confidence in RosettaLigand’s predictive ability for even low-affinity ligands.

## Conclusions

Since the mid-1990’s, successful application of structural information derived from X-ray crystallography and NMR has guided the implementation of computational programs to predict ligand binding^*51*^. Benchmarking algorithms for accurate prediction of protein-ligand binding is highly relevant for improving *in silico* screening of drugs and for structure-guided ligand design. A regular pitfall of these algorithms is the range of errors in predictions for moderate to low-affinity ligands. To that end, our test of RosettaLigand’s ability to dock ligands with affinities in the high micromolar range provides a rigorous test for ligand docking across a range of affinities. The success of Rosetta in predicting the rank order and binding site of these compounds was significant given the few interactions needed for such weak binding. The results here could be used to enhance the algorithm using feedback from NMR on specific binding interactions. Additionally, the identification of accurate binding of low-affinity drug fragments can be used for the designs of higher affinity ligands guided by the context of the binding pocket. Further, as shown here, Rosetta may prove useful for the initial selection of fragments for screening purposes in the development of new drugs for novel protein targets.

## Acknowledgments

This work was supported by NIH grant R01 GM080403 and R01 DA046138. We acknowledge funding by the Deutsche Forschungsgemeinschaft (DFG, German Research Foundation) through SFB1423, project number 421152132, subproject A07. Jens Meiler is further supported by a Humboldt Professorship of the Alexander von Humboldt Foundation. STS received funding through the National Cancer Institute of the National Institutes of Health (F31 CA243353), and the PhRMA Foundation’s Pre-Doctoral Fellowship in Informatics (phrmafoundation.org). This work was conducted using the resources of the Biomolecular NMR facility and the Advanced Computing Center for Research and Education (ACCRE) at Vanderbilt University. We especially would like to acknowledge Dr. Joel Harp for providing assistance in the data collection and analysis of the HisF^C9S^ X-ray crystallography structure. We also especially thank Prof. Markus Voehler for technical assistance with the NMR experiments.

## Author Contributions

Conceptualization, BJB, BA, JM; Data curation, BJB and BA; Formal analysis, BJB and BA; Funding acquisition, JM; Investigation, EO, SS, BJB, BA, SG, AG, XZ; Methodology, BJB and BA; Supervision, KVL and JM.; Writing—original draft, BJB and BA; Writing—review and editing, EO, SS, KVL, JM. All authors have read and agreed to the published version of the manuscript.

## Competing Interests

The authors declare no competing interests.

**Supplemental Table 1:**
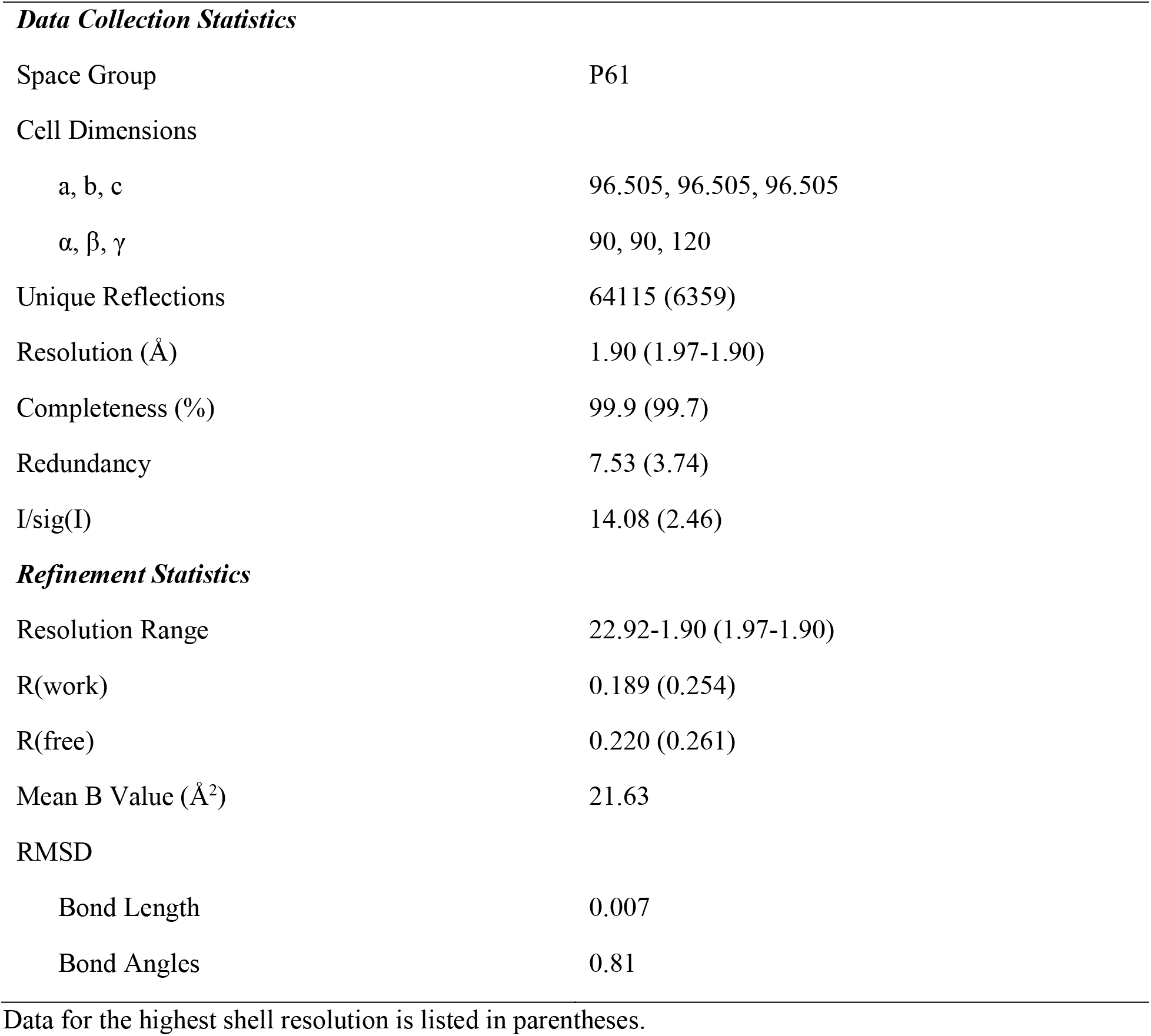
Crystallographic Data Collection for HisF^C9S^.

**Supplementary Figure 1:**
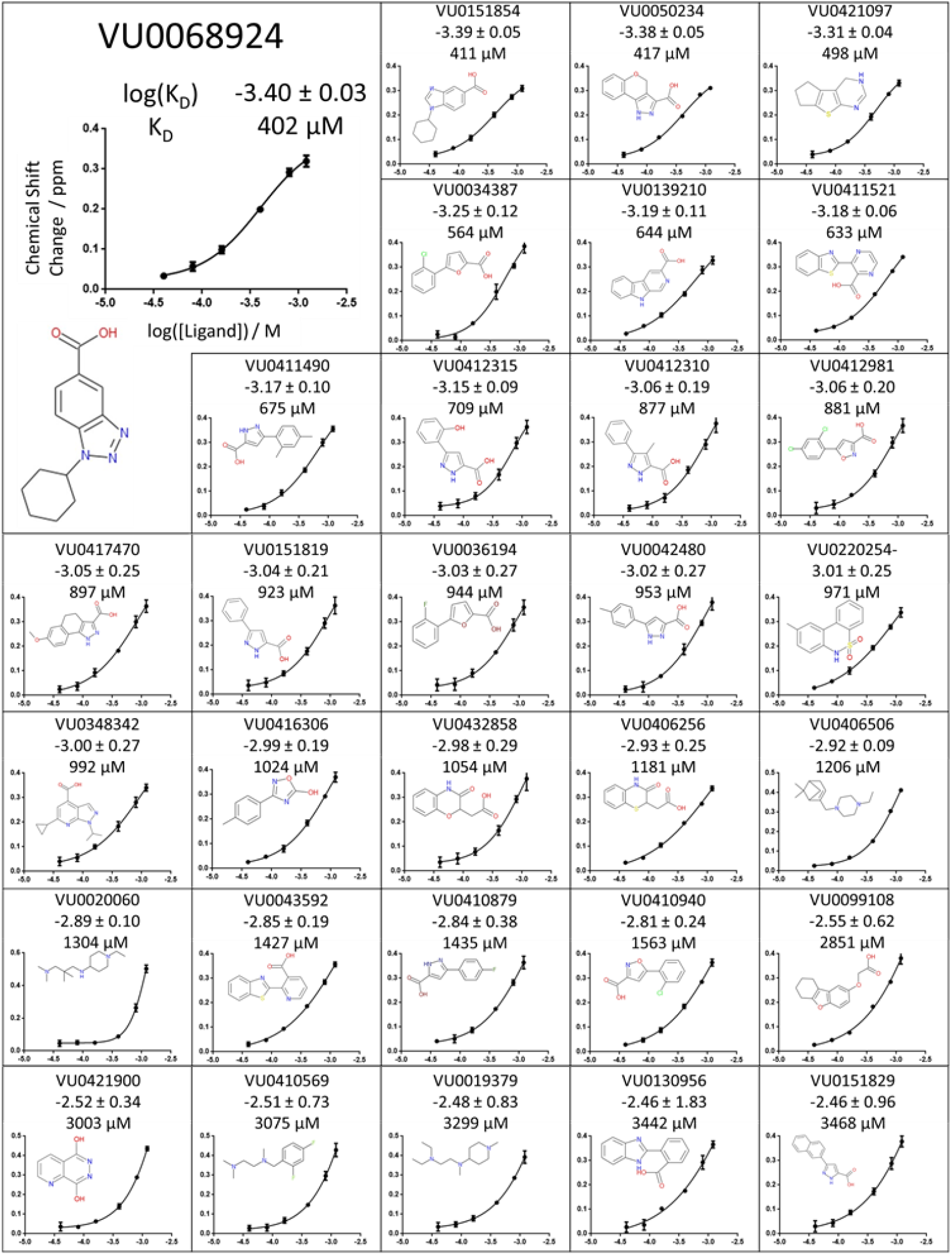
Full dataset for the 31 low-affinity ligands identified for HisF-C9S. For each compound, the 2D structure and binding curve are shown. Also displayed are the log K_D_ and K_D_.

**Supplementary Figure 2:**
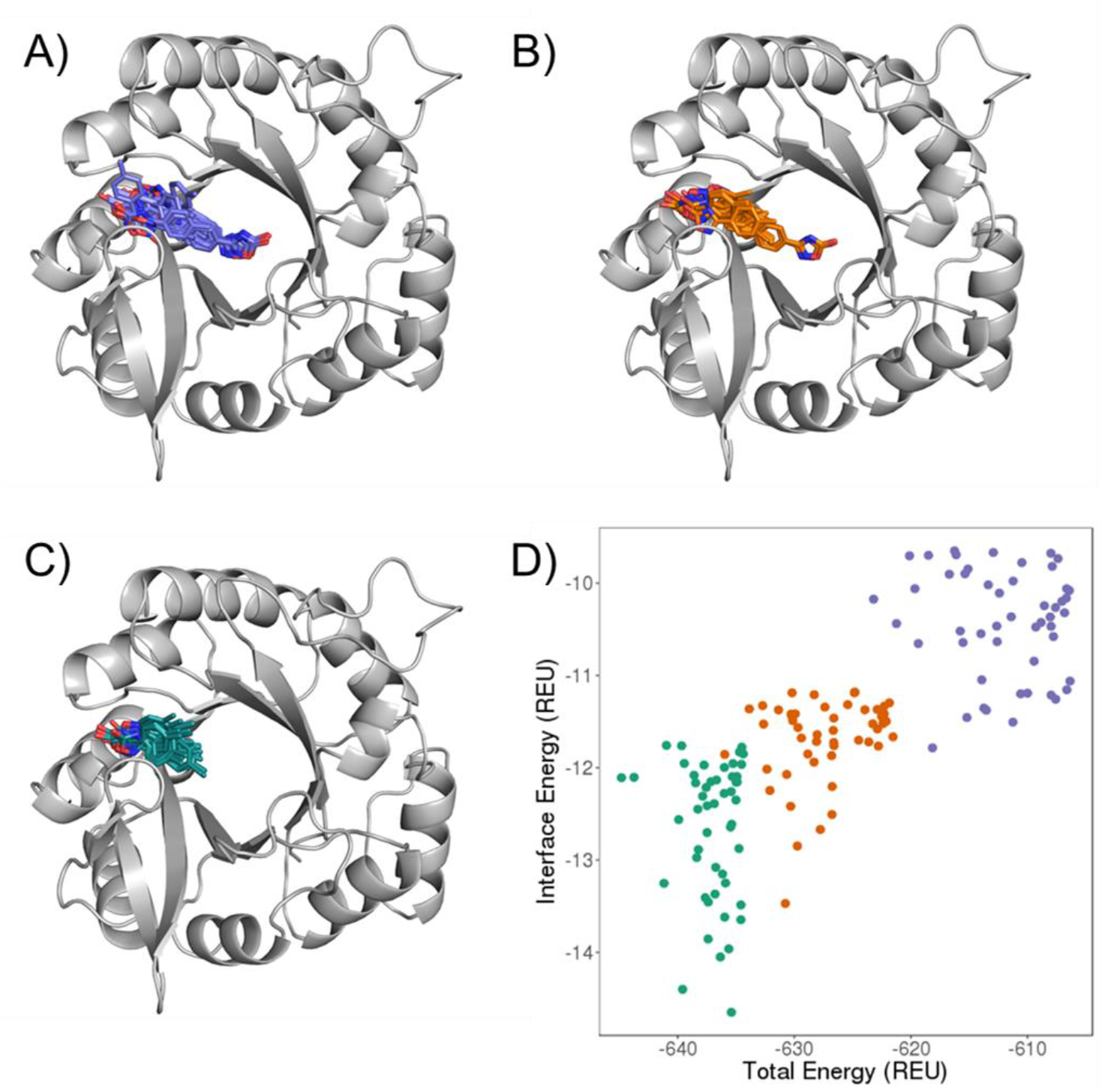
Iterative Docking to Identify Binding Pockets. An example docking strategy in which the results of the first round (**A**) seed docking of a subsequent round (**B**). A final round of docking (**C**) results in a highly converged binding hypothesis. (**D**) Top models from each round were selected based on overall energy and interface binding energy. The values of each docking pose from panels A-C are plotted to show how the energies in both metrics improved with each round of docking.

### Scripts and Commands

Options file, used to specify the input, output, parser file, and packing options. For input files, one must specify the path to the Rosetta database, path to the small molecule PDB file(s), and the path to the small molecule params file. For output, one must specify the type of file to be output (pdb) and number of structures to generate. For the parser, one must specify the path to the XML parser file. For packing, these are standard options to include for side chain repacking.

~~~
-in
      -path
             -database /path_to_database/database/
      -file
             -s /path_to_pdb/filename.pdb
             -extra_res_fa /path_to_params/ligand.params
-out
      -level 300
      -pdb_gz
      -path
            -pdb /path_to_output_files/
            -score /path_to_output_files
      -nstruct 100
      -mute all
      -unmute protocols.jd2.JobDistributor
-parser
      -protocol /path_to_RosettaScripts/RosettaScripts.xml
-packing
      -ex1
      -ex2
      -linmem_ig 10
~~~

XML Script, for the experiments discussed in this study, used to assign values for the cut-off points to detect the protein/small molecule interface, a value for the favor native residue bonus, and values for small molecule translation and rotation. In TASKOPERATIONS:DetectProteinLigandInterface, one must specify values to determine which residues surrounding the small molecule are allowed to be designed and/or repacked (details in my first manuscript), and specify the path to the resfile. In Transform, must specify how much small molecule movement is allowed.

~~~
<ROSETTASCRIPTS>
            <SCOREFXNS>
                      <ligand_soft_rep weights=ligand_soft_rep>
                                      <Reweight scoretype=fa_elec weight=0.42/>
                                      <Reweight scoretype=hbond_bb_sc weight=1.3/>
                                      <Reweight scoretype=hbond_sc weight=1.3/>
                                      <Reweight scoretype=rama weight=0.2/>
                      </ligand_soft_rep>
                      <hard_rep weights=ligandprime>
                                      <Reweight scoretype=fa_intra_rep weight=0.004/>
                                      <Reweight scoretype=fa_elec weight=0.42/>
                                      <Reweight scoretype=hbond_bb_sc weight=1.3/>
                                      <Reweight scoretype=hbond_sc weight=1.3/>
                                      <Reweight scoretype=rama weight=0.2/>
                      </hard_rep>
            </SCOREFXNS>
            <SCORINGGRIDS ligand_chain=“X” width=“16”>
                      <vdw grid_type=“ClassicGrid” weight=“1.0”/>
            </SCORINGGRIDS>
            <TASKOPERATIONS>
                     <DetectProteinLigandInterface name=design_interface cut1=6.0 cut2=8.0 cut3=10.0 cut4=12.0 design=1 resfile=“/path_to_resfile/Resfile_dock”/>
            </TASKOPERATIONS>
            <LIGAND_AREAS>
                     <docking_sidechain chain=X cutoff=6.0 add_nbr_radius=true all_atom_mode=true minimize_ligand=10/>
                     <final_sidechain chain=X cutoff=6.0 add_nbr_radius=true all_atom_mode=true/>
                     <final_backbone chain=X cutoff=7.0 add_nbr_radius=false all_atom_mode=true Calpha_restraints=0.3/>
            </LIGAND_AREAS>
            <INTERFACE_BUILDERS>
                     <side_chain_for_docking ligand_areas=docking_sidechain/>
                     <side_chain_for_final ligand_areas=final_sidechain/>
                     <backbone ligand_areas=final_backbone extension_window=3/>
            </INTERFACE_BUILDERS>
            <MOVEMAP_BUILDERS>
                     <docking sc_interface=side_chain_for_docking minimize_water=true/>
                     <final sc_interface=side_chain_for_final bb_interface=backbone minimize_water=true/>
            </MOVEMAP_BUILDERS>
            <MOVERS>
            single movers
                   <StartFrom name=start_from_X chain=X>
                                <Coordinates x=25.325 y=35.021 z=22.716/>
                   </StartFrom>
                   <FavorNativeResidue name=favor_native bonus=1.0/>
                   <ddG name=calculateDDG jump=1 per_residue_ddg=1 repack=0 scorefxn=hard_rep/>
                   <Transform name=“transform” chain=“X” box_size=“5.0” move_distance=“1.0” angle=“360” cycles=“500” temperature=“5” initial_perturb=“5.0”/>
                   <HighResDocker name=high_res_docker cycles=1 repack_every_Nth=1 scorefxn=ligand_soft_rep movemap_builder=docking/>
                   <PackRotamersMover name=designinterface scorefxn=hard_rep task_operations=design_interface/>
                   <FinalMinimizer name=final scorefxn=hard_rep movemap_builder=final/>
                   <InterfaceScoreCalculator name=add_scores chains=X scorefxn=hard_rep/>
           </MOVERS>
           <PROTOCOLS>
                   <Add mover_name=start_from_X/>
                   <Add mover_name=transform/>
                   <Add mover_name=favor_native/>
                   <Add mover_name=high_res_docker/>
                   <Add mover_name=final/>
                   <Add mover_name=calculateDDG/>
                   <Add mover_name=add_scores/>
          </PROTOCOLS>
</ROSETTASCRIPTS>
~~~

Resfile, used to indicate that residues considered for design and repack are limited to the cut-off points specified above.

#These commands will be applied to all residue positions that lack a specified behavior in the body:

NATAA # allow only native residues (for docking only; no design allowed)

AUTO

start

Rosetta binding site predictions were analyzed using the ddg mover

~~~
<ROSETTASCRIPTS>
          <SCOREFXNS>
                    <hard_rep weights=ligandprime>
                                <Reweight scoretype=fa_intra_rep weight=0.004/>
                                <Reweight scoretype=fa_elec weight=0.42/>
                                <Reweight scoretype=hbond_bb_sc weight=1.3/>
                                <Reweight scoretype=hbond_sc weight=1.3/>
                                <Reweight scoretype=rama weight=0.2/>
                   </hard_rep>
          </SCOREFXNS>
          <MOVERS>
                     <ddG name=calculateDDG jump=1 per_residue_ddg=1 repack_bound=0 repack_unbound=1 scorefxn=hard_rep/>
                     <InterfaceScoreCalculator name=add_scores chains=X scorefxn=hard_rep/>
         </MOVERS>
         <PROTOCOLS>
                     <Add mover_name=calculateDDG/>
                     <Add mover_name=add_scores/>
         </PROTOCOLS>
</ROSETTASCRIPTS>
~~~

## References

[1] Sliwoski, G., Kothiwale, S., Meiler, J., and Lowe, E. W., Jr. (2014) Computational methods in drug discovery, Pharmacol Rev 66, 334–395.

[2] Jain, A. N. (2004) Ligand-based structural hypotheses for virtual screening, J Med Chem 47, 947–961.

[3] Willett, P. (2003) Similarity-based approaches to virtual screening, Biochem Soc Trans 31, 603–606.

[4] Butkiewicz, M., Lowe, E. W., Jr., Mueller, R., Mendenhall, J. L., Teixeira, P. L., Weaver, C. D., and Meiler, J. (2013) Benchmarking ligand-based virtual High-Throughput Screening with the PubChem database, Molecules 18, 735–756.

[5] DeLuca, S., Khar, K., and Meiler, J. (2015) Fully Flexible Docking of Medium Sized Ligand Libraries with RosettaLigand, PLoS One 10, e0132508.

[6] Sousa, S. F., Fernandes, P. A., and Ramos, M. J. (2006) Protein-ligand docking: current status and future challenges, Proteins 65, 15–26.

[7] Rastelli, G., Del Rio, A., Degliesposti, G., and Sgobba, M. (2010) Fast and accurate predictions of binding free energies using MM-PBSA and MM-GBSA, Journal of computational chemistry 31, 797–810.

[8] Masetti, M., Cavalli, A., Recanatini, M., and Gervasio, F. L. (2009) Exploring complex protein-ligand recognition mechanisms with coarse metadynamics, The journal of physical chemistry. B 113, 4807–4816.

[9] Cozza, G., Bonvini, P., Zorzi, E., Poletto, G., Pagano, M. A., Sarno, S., Donella-Deana, A., Zagotto, G., Rosolen, A., Pinna, L. A., Meggio, F., and Moro, S. (2006) Identification of ellagic acid as potent inhibitor of protein kinase CK2: a successful example of a virtual screening application, J Med Chem 49, 2363–2366.

[10] Becker, O. M., Dhanoa, D. S., Marantz, Y., Chen, D., Shacham, S., Cheruku, S., Heifetz, A., Mohanty, P., Fichman, M., Sharadendu, A., Nudelman, R., Kauffman, M., and Noiman, S. (2006) An integrated in silico 3D model-driven discovery of a novel, potent, and selective amidosulfonamide 5-HT1A agonist (PRX-00023) for the treatment of anxiety and depression, J Med Chem 49, 3116–3135.

[11] Durrant, J. D., Hall, L., Swift, R. V., Landon, M., Schnaufer, A., and Amaro, R. E. (2010) Novel naphthalene-based inhibitors of Trypanosoma brucei RNA editing ligase 1, PLoS neglected tropical diseases 4, e803.

[12] Schueler-Furman, O., Wang, C., Bradley, P., Misura, K., and Baker, D. (2005) Progress in modeling of protein structures and interactions, Science 310, 638–642.

[13] Leaver-Fay, A., Tyka, M., Lewis, S. M., Lange, O. F., Thompson, J., Jacak, R., Kaufman, K., Renfrew, P. D., Smith, C. A., Sheffler, W., Davis, I. W., Cooper, S., Treuille, A., Mandell, D. J., Richter, F., Ban, Y. E., Fleishman, S. J., Corn, J. E., Kim, D. E., Lyskov, S., Berrondo, M., Mentzer, S., Popovic, Z., Havranek, J. J., Karanicolas, J., Das, R., Meiler, J., Kortemme, T., Gray, J. J., Kuhlman, B., Baker, D., and Bradley, P. (2011) ROSETTA3: an object-oriented software suite for the simulation and design of macromolecules, Methods in enzymology 487, 545–574.

[14] Meiler, J., and Baker, D. (2006) ROSETTALIGAND: protein-small molecule docking with full side-chain flexibility, Proteins 65, 538–548.

[15] Davis, I. W., and Baker, D. (2009) RosettaLigand docking with full ligand and receptor flexibility, J Mol Biol 385, 381–392.

[16] Lemmon, G., Kaufmann, K., and Meiler, J. (2012) Prediction of HIV-1 protease/inhibitor affinity using RosettaLigand, Chemical biology & drug design 79, 888–896.

[17] Kaufmann, K. W., Dawson, E. S., Henry, L. K., Field, J. R., Blakely, R. D., and Meiler, J. (2009) Structural determinants of species-selective substrate recognition in human and Drosophila serotonin transporters revealed through computational docking studies, Proteins 74, 630–642.

[18] Nguyen, E. D., Norn, C., Frimurer, T. M., and Meiler, J. (2013) Assessment and challenges of ligand docking into comparative models of G-protein coupled receptors, PLoS One 8, e67302.

[19] Combs, S. A., Deluca, S. L., Deluca, S. H., Lemmon, G. H., Nannemann, D. P., Nguyen, E. D., Willis, J. R., Sheehan, J. H., and Meiler, J. (2013) Small-molecule ligand docking into comparative models with Rosetta, Nat Protoc 8, 1277–1298.

[20] Kaufmann, K. W., and Meiler, J. (2012) Using RosettaLigand for small molecule docking into comparative models, PLoS One 7, e50769.

[21] Kufareva, I., Katritch, V., Participants of, G. D., Stevens, R. C., and Abagyan, R. (2014) Advances in GPCR modeling evaluated by the GPCR Dock 2013 assessment: meeting new challenges, Structure 22, 1120–1139.

[22] Jiang, L., Althoff, E. A., Clemente, F. R., Doyle, L., Rothlisberger, D., Zanghellini, A., Gallaher, J. L., Betker, J. L., Tanaka, F., Barbas, C. F., 3rd, Hilvert, D., Houk, K. N., Stoddard, B. L., and Baker, D. (2008) De novo computational design of retro-aldol enzymes, Science 319, 1387–1391.

[23] Procko, E., Hedman, R., Hamilton, K., Seetharaman, J., Fleishman, S. J., Su, M., Aramini, J., Kornhaber, G., Hunt, J. F., Tong, L., Montelione, G. T., and Baker, D. (2013) Computational design of a protein-based enzyme inhibitor, Journal of molecular biology 425, 3563–3575.

[24] Rothlisberger, D., Khersonsky, O., Wollacott, A. M., Jiang, L., DeChancie, J., Betker, J., Gallaher, J. L., Althoff, E. A., Zanghellini, A., Dym, O., Albeck, S., Houk, K. N., Tawfik, D. S., and Baker, D. (2008) Kemp elimination catalysts by computational enzyme design, Nature 453, 190–195.

[25] Wang, L., Althoff, E. A., Bolduc, J., Jiang, L., Moody, J., Lassila, J. K., Giger, L., Hilvert, D., Stoddard, B., and Baker, D. (2012) Structural analyses of covalent enzyme-substrate analog complexes reveal strengths and limitations of de novo enzyme design, Journal of molecular biology 415, 615–625.

[26] Price, A. J., Howard, S., and Cons, B. D. (2017) Fragment-based drug discovery and its application to challenging drug targets, Essays in biochemistry 61, 475–484.

[27] de Kloe, G. E., Bailey, D., Leurs, R., and de Esch, I. J. (2009) Transforming fragments into candidates: small becomes big in medicinal chemistry, Drug discovery today 14, 630–646.

[28] Siegal, G., Ab, E., and Schultz, J. (2007) Integration of fragment screening and library design, Drug discovery today 12, 1032–1039.

[29] Jhoti, H., Williams, G., Rees, D. C., and Murray, C. W. (2013) The ‘rule of three’ for fragment-based drug discovery: where are we now?, Nature reviews. Drug discovery 12, 644–645.

[30] Shuker, S. B., Hajduk, P. J., Meadows, R. P., and Fesik, S. W. (1996) Discovering high-affinity ligands for proteins: SAR by NMR, Science 274, 1531–1534.

[31] Hajduk, P. J., Gerfin, T., Boehlen, J. M., Haberli, M., Marek, D., and Fesik, S. W. (1999) High-throughput nuclear magnetic resonance-based screening, J Med Chem 42, 2315–2317.

[32] Bender, B. J., Cisneros, A., 3rd, Duran, A. M., Finn, J. A., Fu, D., Lokits, A. D., Mueller, B. K., Sangha, A. K., Sauer, M. F., Sevy, A. M., Sliwoski, G., Sheehan, J. H., DiMaio, F., Meiler, J., and Moretti, R. (2016) Protocols for Molecular Modeling with Rosetta3 and RosettaScripts, Biochemistry 55, 4748–4763.

[33] Levi, R. N., and Waxman, S. (1975) Schizophrenia, epilepsy, cancer, methionine, and folate metabolism. Pathogenesis of schizophrenia, Lancet 2, 11–13.

[34] Lipchock, J. M., and Loria, J. P. (2008) 1H, 15N and 13C resonance assignment of imidazole glycerol phosphate (IGP) synthase protein HisF from Thermotoga maritima, Biomol NMR Assign 2, 219–221.

[35] Schanda, P., and Brutscher, B. (2005) Very fast two-dimensional NMR spectroscopy for real-time investigation of dynamic events in proteins on the time scale of seconds, J Am Chem Soc 127, 8014–8015.

[36] Delaglio, F., Grzesiek, S., Vuister, G. W., Zhu, G., Pfeifer, J., and Bax, A. (1995) NMRPipe: a multidimensional spectral processing system based on UNIX pipes, J Biomol NMR 6, 277–293.

[37] Lee, W., Tonelli, M., and Markley, J. L. (2015) NMRFAM-SPARKY: enhanced software for biomolecular NMR spectroscopy, Bioinformatics 31, 1325–1327.

[38] Wu, B., Zhang, Z., Noberini, R., Barile, E., Giulianotti, M., Pinilla, C., Houghten, R. A., Pasquale, E. B., and Pellecchia, M. (2013) HTS by NMR of combinatorial libraries: a fragment-based approach to ligand discovery, Chem Biol 20, 19–33.

[39] Farmer, B. T., 2nd, Constantine, K. L., Goldfarb, V., Friedrichs, M. S., Wittekind, M., Yanchunas, J., Jr., Robertson, J. G., and Mueller, L. (1996) Localizing the NADP+ binding site on the MurB enzyme by NMR, Nat Struct Biol 3, 995–997.

[40] Fielding, L. (2003) NMR methods for the determination of protein-ligand dissociation constants, Curr Top Med Chem 3, 39–53.

[41] Lemmon, G., and Meiler, J. (2012) Rosetta Ligand docking with flexible XML protocols, Methods in molecular biology 819, 143–155.

[42] Fleishman, S. J., Leaver-Fay, A., Corn, J. E., Strauch, E. M., Khare, S. D., Koga, N., Ashworth, J., Murphy, P., Richter, F., Lemmon, G., Meiler, J., and Baker, D. (2011) RosettaScripts: a scripting language interface to the Rosetta macromolecular modeling suite, PLoS One 6, e20161.

[43] Sing, T., Sander, O., Beerenwinkel, N., and Lengauer, T. (2005) ROCR: visualizing classifier performance in R, Bioinformatics 21, 3940–3941.

[44] Lang, D., Thoma, R., Henn-Sax, M., Sterner, R., and Wilmanns, M. (2000) Structural evidence for evolution of the beta/alpha barrel scaffold by gene duplication and fusion, Science 289, 1546–1550.

[45] Harner, M. J., Frank, A. O., and Fesik, S. W. (2013) Fragment-based drug discovery using NMR spectroscopy, J Biomol NMR 56, 65–75.

[46] Congreve, M., Carr, R., Murray, C., and Jhoti, H. (2003) A ‘rule of three’ for fragment-based lead discovery?, Drug discovery today 8, 876–877.

[47] Chaudhuri, B. N., Lange, S. C., Myers, R. S., Davisson, V. J., and Smith, J. L. (2003) Toward understanding the mechanism of the complex cyclization reaction catalyzed by imidazole glycerolphosphate synthase: crystal structures of a ternary complex and the free enzyme, Biochemistry 42, 7003–7012.

[48] Su, M., Yang, Q., Du, Y., Feng, G., Liu, Z., Li, Y., and Wang, R. (2019) Comparative Assessment of Scoring Functions: The CASF-2016 Update, J Chem Inf Model 59, 895–913.

[49] Williamson, M. P. (2013) Using chemical shift perturbation to characterise ligand binding, Prog Nucl Magn Reson Spectrosc 73, 1–16.

[50] Fu, D. Y., and Meiler, J. (2018) Predictive Power of Different Types of Experimental Restraints in Small Molecule Docking: A Review, Journal of chemical information and modeling 58, 225–233.

[51] Waszkowycz, B., Clark, D. E., and Gancia, E. (2011) Outstanding challenges in protein-ligand docking and structure-based virtual screening, In Wiley Interdisciplinary Reviews: Computational Molecular Science, pp 229–259.

